# Dynamic Maternal Gradients Control Timing and Shift-Rates for *Drosophila* Gap Gene Expression

**DOI:** 10.1101/068064

**Authors:** Berta Verd, Anton Crombach, Johannes Jaeger

## Abstract

Pattern formation during development is a highly dynamic process. In spite of this, few experimental and modelling approaches take into account the explicit time-dependence of the rules governing regulatory systems. We address this problem by studying dynamic morphogen interpretation by the gap gene network in *Drosophila melanogaster*. Gap genes are involved in segment determination during early embryogenesis. They are activated by maternal morphogen gradients encoded by *bicoid (bcd)* and *caudal (cad)*. These gradients decay at the same time-scale as the establishment of the antero-posterior gap gene pattern. We use a reverse-engineering approach, based on data-driven regulatory models called gene circuits, to isolate and characterise the explicitly time-dependent effects of changing morphogen concentrations on gap gene regulation. To achieve this, we simulate the system in the presence and absence of dynamic gradient decay. Comparison between these simulations reveals that maternal morphogen decay controls the timing and limits the rate of gap gene expression. In the anterior of the embyro, it affects peak expression and leads to the establishment of smooth spatial boundaries between gap domains. In the posterior of the embryo, it causes a progressive slow-down in the rate of gap domain shifts, which is necessary to correctly position domain boundaries and to stabilise the spatial gap gene expression pattern. We use a newly developed method for the analysis of transient dynamics in non-autonomous (time-variable) systems to understand the regulatory causes of these effects. By providing a rigorous mechanistic explanation for the role of maternal gradient decay in gap gene regulation, our study demonstrates that such analyses are feasible and reveal important aspects of dynamic gene regulation which would have been missed by a traditional steady-state approach. More generally, it highlights the importance of transient dynamics for understanding complex regulatory processes in development.

**Author Summary:** Animal development is a highly dynamic process. Biochemical or environmental signals can cause the rules that shape it to change over time. We know little about the effects of such changes. For the sake of simplicity, we usually leave them out of our models and experimental assays. Here, we do exactly the opposite. We characterise precisely those aspects of pattern formation caused by changing signalling inputs to a gene regulatory network, the gap gene system of *Drosophila melanogaster*. Gap genes are involved in determining the body segments of flies and other insects during early development. Gradients of maternal morphogens activate the expression of the gap genes. These gradients are highly dynamic themselves, as they decay while being read out. We show that this decay controls the peak concentration of gap gene products, produces smooth boundaries of gene expression, and slows down the observed positional shifts of gap domains in the posterior of the embryo, thereby stabilising the spatial pattern. Our analysis demonstrates that the dynamics of gene regulation not only affect the timing, but also the positioning of gene expression. This suggests that we must pay closer attention to transient dynamic aspects of development than is currently the case.

## Introduction

Biological systems depend on time. Like everything else that persists for more than an instant, there is a temporal dimension to their existence. This much is obvious. What is less obvious, however, is the active role that time plays in altering the rules governing biological processes. For instance, fluctuating environmental conditions modify the selective pressures that drive adaptive evolutionary change [1,3–5], time-dependent inductive signals or environmental cues trigger and remodel developmental pathways [6,7], and dynamic morphogen gradients influence patterning, not only across space but also through time [8–16]. In spite of this, many current attempts at understanding biological processes neglect important aspects of this temporal dimension [17]. For practical reasons, experimental studies often glance over the detailed dynamics of a process, and focus on its end product or output pattern instead. Similarly, modelling studies frequently restrict themselves to a small-enough time window allowing them to ignore temporal changes in the rules governing the system. Accuracy is sacrificed and the scope of the investigation limited for the sake of simplicity and tractability. Although reasonable, and often even necessary, such simplifications can lead us to miss important aspects of biological regulatory dynamics.

We set out to tackle explicitly time-dependent aspects of morphogen interpretation for pattern formation during animal development. As a case study, we use the gap gene network, which is involved in segment determination during the blastoderm stage of early development in the vinegar fly *Drosophila melanogaster* [18]. Activated by long-range gradients of maternal morphogens Bicoid (Bcd) and Caudal (Cad), the trunk gap genes *hunchback (hb)*, *Krüppel (Kr)*, *giant (gt)*, and *knirps (kni)* become expressed in broad overlapping domains along the antero-posterior (A–P) axis of the embryo (Fig. 1). The establishment of these domains is fast and dynamic. Subsequently, gap gene domain boundaries sharpen and domains in the posterior region of the embryo shift anteriorly over time (Fig. 1). Towards the end of the blastoderm stage, gap gene production rates drop and domain shifts slow down. The blastoderm stage ends with the onset of gastrulation.

**Figure 1.**
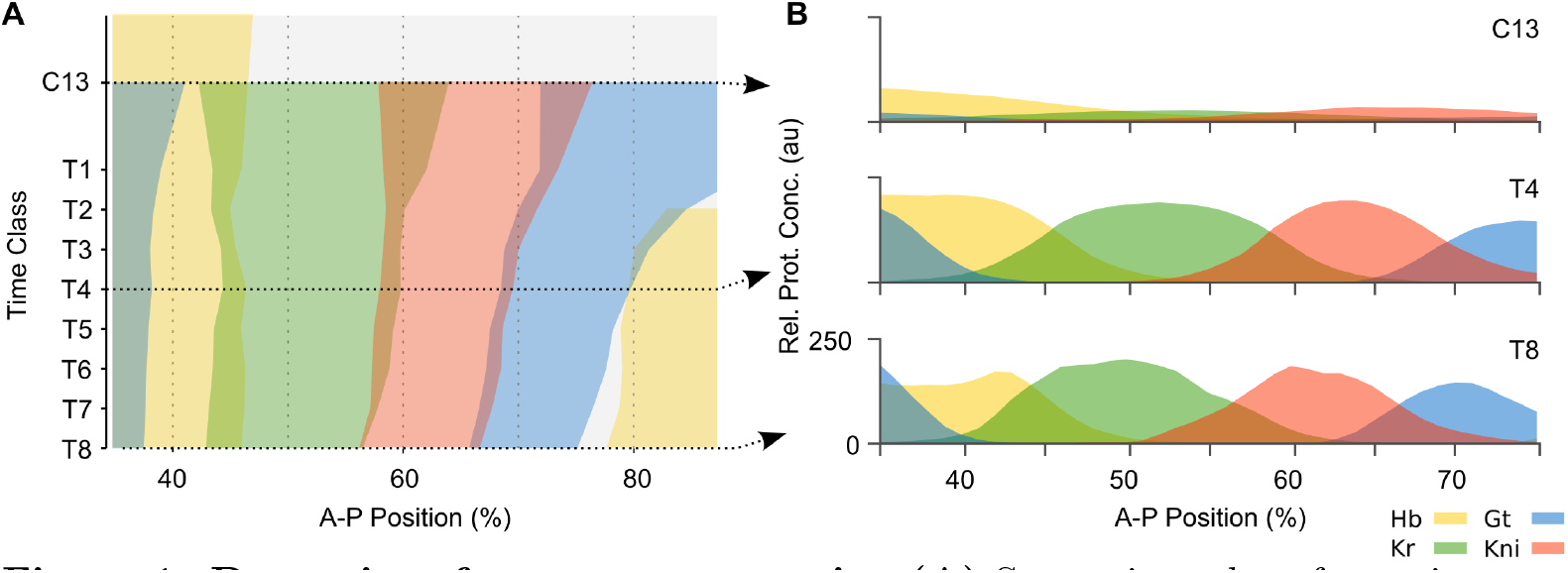
Dynamics of gap gene expression. **(A)** Space-time plot of protein expression data for the trunk gap genes during the late blastoderm stage in *D. melanogaster*. Coloured areas demarcate regions with relative protein concentration above half-maximum value. Time flows downwards along the y-axis. **(B)** Cross-sections of gene expression in (A) at cycle C13 and time classes C14A-T4 and T8 (dashed arrows in (A)). Y-axes indicate relative protein concentration in arbitrary units (au). In both panels, x-axes represent %A–P position, where 0% is the anterior pole. Hunchback (Hb) is shown in yellow, Krüppel (Kr) in green, Knirps (Kni) in red, and Giant (Gt) in blue. C13: cleavage cycle 13; C14A-T1–8: cleavage cycle 14A, time classes 1–8 (see Models and Methods for details).

The gap gene system is one of the most thoroughly studied developmental gene regulatory networks today. For our particular purposes, we take advantage of the fact that it has been extensively reverse-engineered using data-driven modelling. This approach is based on fitting dynamical models of gap gene regulation, called gene circuits, to quantitative spatio-temporal gene expression data [19–27,29,34,35].

Dynamical models capture how a given regulatory process unfolds over time. They are frequently formulated in terms of ordinary differential equations (ODEs) with parameter values that remain constant over time. Such equations represent an *autonomous dynamical system*. Central to the analysis of such dynamical systems is the concept of phase space and its associated features (S1 Figure A). Phase (or state) space is an abstract space that contains all possible states of a system. Its axes are defined by the state variables, which in our case represent the concentrations of transcription factors encoded by the gap genes. Trajectories through phase space describe how a system's state changes as time progresses. The trajectories of a gap gene circuit describe how transcription factor concentrations change over time. All trajectories taken together constitute the flow of the system. This flow is shaped by the regulatory structure of the underlying network—the type (activation/repression) and strength of interactions between the constituent factors—which is given by the system's parameters. Since these parameters are constant over time in an autonomous system, the trajectories are fully determined given a specific set of initial conditions. Once the system's variables no longer change, it has reached a steady state. Steady states can be stable—such as attractors with converging trajectories from all directions defining a basin of attraction—or unstable—such as saddles; where trajectories converge only along certain directions and diverge along others. The type and arrangement of steady states, and their associated basins of attraction define the *phase portrait* of the system (S1 Figure A). There exist powerful analytical tools to analyse and understand the phase portrait and the range of dynamic behaviours determined by it. Geometrical analysis of the phase portrait enables us to build up a rich qualitative understanding of the dynamics of non-linear autonomous systems without solving the underlying equations analytically [36].

The application of dynamical systems concepts and phase space analysis to the study of cellular and developmental processes has a long history (see [37–39] for recent reviews). In particular, it has been successfully applied to the study of the gap gene system. Manu and colleagues [22,23,40] examined the dynamics and robustness of gap gene regulation in *D. melanogaster* using diffusion-less gene circuits fit to quantitative expression data. These models have a four-dimensional phase space, where the axes represent the concentrations of transcription factors encoded by the trunk gap genes *hb*, *Kr*, *gt*, and *kni*. The analysis of these phase portraits yields a rigorous understanding of the patterning capabilities of the system.

The analysis by Manu *et al.* [23] corroborated and expanded upon earlier genetic evidence [41] indicating that the regulatory dynamics responsible for domain boundary placement in the anterior versus the posterior of the embryo are very different. In the anterior, spatial boundaries of gap gene expression domains are positioned statically, meaning that they remain in place over time [42]. Stationary boundaries are regulated in two distinct ways [23]. (1) In the case of the posterior boundary of the anterior *gt* domain, different nuclei along the A–P axis have equivalent attractors positioned at different locations in phase space (shift in attractor position); (2) in the case of the posterior boundary of the anterior *hb* domain, system trajectories fall into different basins of attraction (attractor selection) (Fig. 2A). In both of these cases, patterning is largely governed by the position of attractors in a multi-stable phase space.

In contrast, gap domain boundaries in the posterior of the embryo shift anteriorly over time [25,42]. In this region, the system always remains far from steady state, and the dynamics of gene expression are transient. Therefore, trajectories here are fairly independent of precise attractor positions. The model by Manu *et al.* [23] shows that posterior gap gene expression is governed by an unstable manifold (Fig. 2A). An unstable manifold is the trajectory connecting a saddle to an attractor (S1 Figure A). The authors demonstrate that this manifold has canalising properties since it compresses many incoming neighbouring trajectories into an increasingly smaller sub-volume of phase space over time [23]. This explains the observed robustness of posterior patterning. Moreover, the geometry of the unstable manifold provides an explanation for the ordered succession of gap genes that become expressed in each nucleus of the posterior region. Such an ordered temporal sequence of gene expression, if arranged appropriately along the A–P axis, creates the observed kinematic anterior shifts of gap domains over time (Fig. 2A).

**Figure 2.**
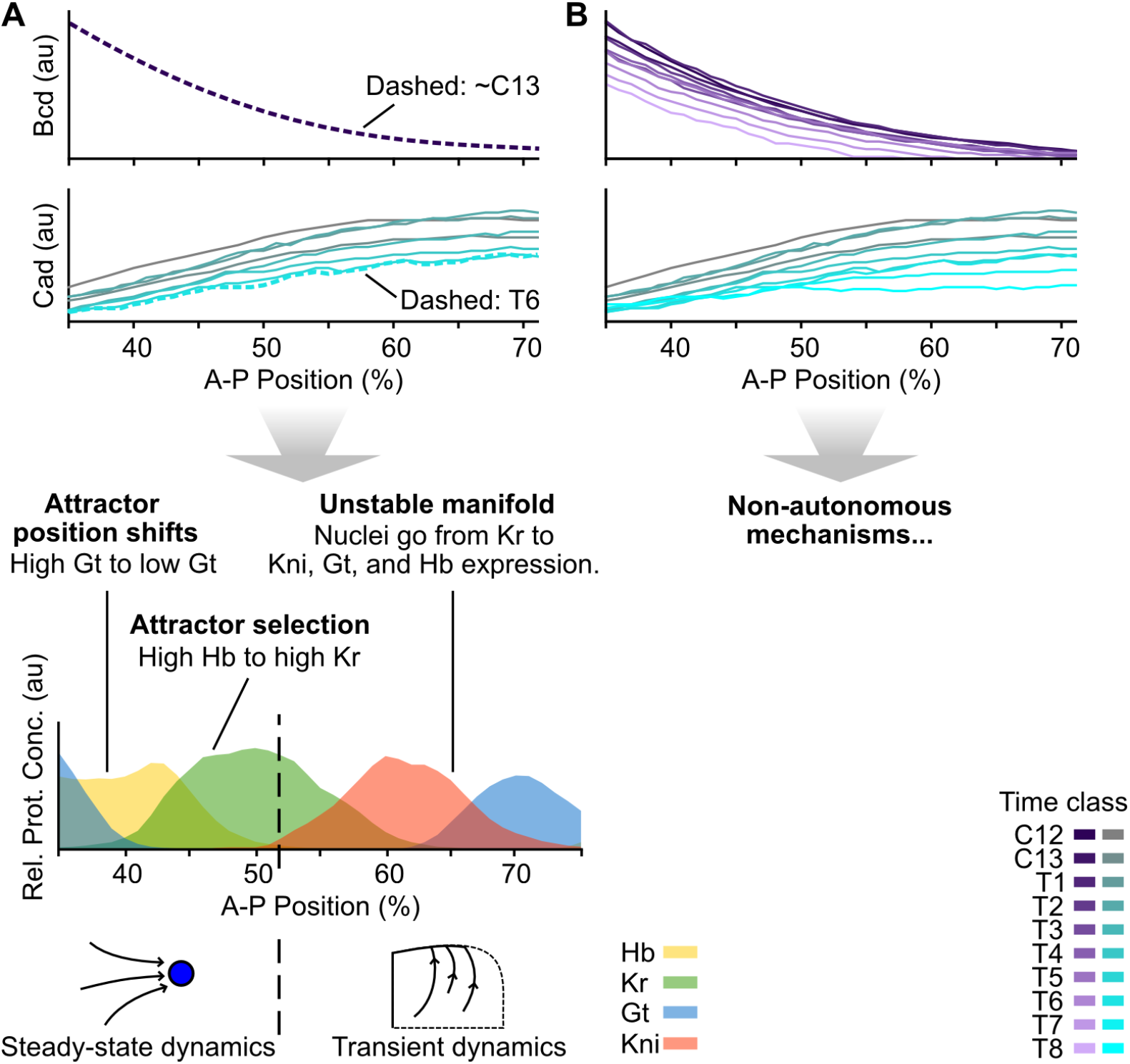
Static-Bcd versus non-autonomous gap gene patterning mechanisms. **(A)** Summary of the phase space analysis of the Static-Bcd gap gene circuit in *D. melanogaster* by Manu *et. al.* [23]. An exponential function fit to the Bcd profile at cycle C13 was used to calculate trajectories and phase portraits. All Cad profiles until time class T6 were considered for simulating trajectories, but phase portraits were calculated using the profile at T6 only. This gene circuit displays the following mechanisms for boundary formation: patterning between 35–51% A–P position takes place in a multi-stable regime close to steady state. The Gt boundary is established as the relevant attractor moves from high to low Gt concentration in more posterior nuclei. The Hb-Kr interface forms as the maternal Hb gradient places more anterior nuclei in the basin of an attractor with high Hb concentration, and more posterior nuclei in the basin of an attractor at high Kr concentration. Between 51 and 53% A–P position a saddle-node bifurcation takes place, and the dynamics become transient in nuclei posterior of 52%. These nuclei are all in the basin of the same attractor and approach it by first converging towards an unstable manifold. Anterior shifts in these posterior gap gene domains emerge from a coordinated succession of trajectories in more posterior nuclei approaching the unstable manifold. See [23] for details. **(B)** In the non-autonomous gap gene circuit analysed here, Bcd and Cad gradient profiles are included for every time point. They are used to calculate trajectories of the system and instantaneous phase portraits as discussed in the main text. Plots in (A) and (B) show % A–P position along the x-axes, and protein concentrations (in arbitrary units, au) along the y-axes as in Fig.1B).

Despite its explanatory power, the analysis by Manu *et al.* [23] is limited in an important way. In order to simplify phase space analysis, the authors implement simplified dynamics of maternal morphogens Bcd and Cad in their model (Fig. 2A). They use a time-invariant exponential approximation to simulate the Bcd gradient and Cad is assumed to reach a steady-state profile about 20-30 minutes before gastrulation [22, 23]. This steady-state profile is used for model analysis. (Based on this, we will refer to this formulation as the static-Bcd gene circuit model in what follows). Although reasonable, these simplifications affect the accuracy of the model, since Bcd and Cad have their own expression dynamics on a similar time scale as gap proteins. The Bcd gradient decays and Cad clears from much of the posterior trunk region towards the end of the blastoderm stage (Fig. 2B) [42]. This means that the autonomous analysis of the static-Bcd model is not well suited to investigate the dynamic interpretation of morphogen gradients. In particular, assuming autonomy makes it impossible to isolate and study the explicitly time-dependent effects of changing gradient concentrations on gap gene regulation and pattern formation.

For this reason, we consider the dynamics of maternal morphogens explicitly in our model. We have obtained gap gene circuits that incorporate realistic time-variable maternal gradients of Bcd and Cad (Fig. 2B) [26]. These gradients are implemented as external inputs to gap gene regulation (see Models and Methods section). They are not influenced by any of the state variables and, thus, are parameters of the system. This means that our gap gene circuits become fully *non-autonomous* [54], since certain parameter values now change over time. While non-autonomous equations are not significantly more difficult to formulate or simulate than autonomous ones, phase space analysis is far from trivial. As model parameters change, so does the geometry of the phase portrait, and consequently system trajectories are actively shaped by this time-dependence. Separatrices and attractors can change their position (*geometrical change*), and steady states can be created and annihilated through *bifurcation* events (*topological change*) (S1 Figure B). In autonomous systems, bifurcations can only occur along the spatial axis of the model. In non-autonomous systems, they also occur in time, implying that trajectories can switch from one basin of attraction to another during a simulation run. We can think of time-variable phase portraits as embedded in *parameter-space*. We call the combination of phase and parameter space the *configuration space* of the system. The configuration space on non-autonomous models hence encodes a much richer repertoire of dynamical mechanisms of pattern formation than autonomous phase space alone. This can complicate analysis and interpretation of the system considerably.

Using a simple model of a genetic toggle switch, we have established a methodology for the characterisation of transient dynamics in non-autonomous systems (S1 Figure B), based on the analysis of *instantaneous phase portraits* [43, 45]. Such portraits are generated by fixing the values of system parameters starting at a given point in time, and then determining the geometrical arrangement of saddles, attractors, and their basins under these “frozen” conditions. The overall non-autonomous trajectory of the system is given by a series of instantaneous phase portraits over time. With sufficiently high temporal resolution, this method yields an accurate picture of the non-autonomous mechanisms of pattern formation implemented by the system. These mechanisms can be classified into four broad categories [43]: (1) *transitions* of the system from one steady state to another, (2) pursuit of a moving attractor within a basin of attraction, (3) *geometrical capture*, where a trajectory crosses a separatrix, and (4) *topological capture*, where a trajectory suddenly falls into a new basin of attraction due to a preceding bifurcation event (S1 Figure B). This classification scheme can be used to characterise the dynamical repertoire of non-autonomous models in a way analogous to phase space analysis in autonomous dynamical systems.

In this paper, we present a detailed analysis of a non-autonomous gap gene circuit. Specifically, we use the model to address the effect of non-autonomy, *i. e.* the effect of time-variable maternal gradient concentrations, on gap gene regulation (Fig. 2). To isolate explicitly time-dependent regulatory aspects, we simulate gap gene expression in the presence and absence of maternal gradient decay. Using phase space analysis, we then identify and characterise the dynamic regulatory mechanisms responsible for the observed differences between the two simulations. Our analysis reveals that maternal gradient decay limits the levels of gap gene expression and controls the dynamical positioning of posterior domains by regulating the rate and timing of domain shifts in the posterior of the embryo.

## Models and Methods

### Non-autonomous gene circuits

Non-autonomous gene circuit models are based on the connectionist formalism introduced by Mjolsness *et al.* [21], modified to include time-variable external regulatory inputs as previously described [26,34]. Gene circuits are hybrid models with discrete cell divisions and continuous gene regulatory dynamics. The basic objects of the model consist of nuclei arranged in a one-dimensional row along the A–P axis of the embryo, covering the trunk region between 35 and 92% A–P position (where 0% is the anterior pole). Models include the last two cleavage cycles of the blastoderm stage (C13 and C14A) and end with the onset of gastrulation; C14A is further subdivided into eight time classes of equal duration (T1–T8). At the end of C13, division occurs and the number of nuclei doubles.

The state variables of the system consist of the concentration levels of proteins produced by the trunk gap genes *hb*, *Kr*, *gt*, and *kni*. We denote the concentration of gap protein *a* in nucleus *i* at time *t* by 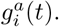 Change in protein concentration over time is given by the following set of ODEs:

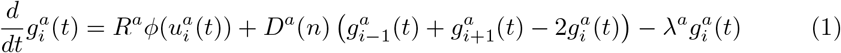

where *R^a^*, *D^a^* and *λ^a^* are rates of protein production, diffusion, and decay, respectively. Diffusion depends on the distance between neighbouring nuclei, which halves at nuclear division; thus, *D^a^* depends on the number of preceding divisions *n*. *ϕ* is a sigmoid regulation-expression function representing coarse-grained kinetics of transcriptional regulation. It is defined as follows:

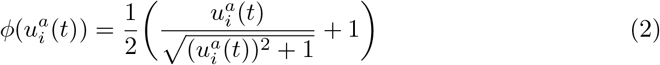

where

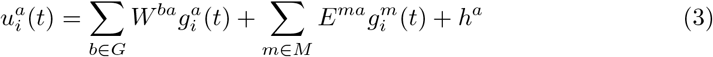

with the set of trunk gap genes *G* = {*hb*, *Kr*, *gt*, *kni*}, and the set of external regulatory inputs *M* = {Bcd, Cad, Tll, Hkb}. External regulator concentrations 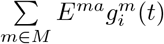 are interpolated from quantified spatio-temporal protein expression profiles [26, 42, 46]. The dynamic nature of these profiles renders the parameter term representing external regulatory inputs 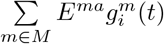 time-dependent; explicit time-dependence of parameters implies non-autonomy of the dynamical system (see Introduction and [54]).

Interconnectivity matrices *W* and *E* define interactions among gap genes, as well as regulatory inputs from external inputs, respectively. The elements of these matrices, *w^ba^* and *e^ma^*, are called regulatory weights. They encode the effect of regulator *b* or *m* on target gene *a*. These weights may be positive (representing an activating regulatory input), negative (representing repression), or near zero (representing the absence of a regulatory interaction). *h^a^* is a threshold parameter that represents the activation state of target gene a in the absence of any spatially and temporally specific regulatory input. This term incorporates the regualtory influence of factors that are not expressed in a spatially specific manner (for example, the pioneer factor Zelda [31]). Equation (1) determines regulatory dynamics during interphase. In order to accurately implement the non-instantaneous duration of the nuclear division between C13 and C14A, the production rate *R^a^* is set to zero during a mitotic phase, which immediately precedes the instantaneous nuclear division. Mitotic schedule as in [26].

### Model fitting and selection

We determine the values for parameters *R^a^*, *λ^a^*, *W*, *E*, and *h^a^* using a reverse-engineering approach [19, 25, 26, 34]. For this purpose, we numerically solve gene circuit equations (1) across the region between 35 and 92% A–P position using a Runge-Kutta Cash-Karp adaptive step-size solver [26]. Models are fit to a previously published quantitative data set of spatio-temporal gap protein expression [26, 42, 46] (see Fig. 1 for gap gene expression patterns, and Fig. 2B for dynamic Bcd and Cad profiles). Model fitting was performed using a global optimization algorithm called parallel Lam Simulated Annealing (pLSA) [47]. We use a weighted least squares cost function as previously described [26].

To enable comparison of our results to the static-Bcd gene circuit analysis by Manu *et al.* [23], we keep model formalism and fitting procedure as similar as possible to this earlier study. Manu and colleagues fitted gene circuits including a diffusion term, but analysed the model with diffusion rates *D^a^* set to zero [23]. This diffusion-less approach reduces the phase space of the model from hundreds of dimensions to 4 by spatially uncoupling the equations and considering each nucleus independently from its neighbours. Dimensionality reduction is essential for geometrical analysis of phase space. Unfortunately, setting diffusion to zero in our best 3 (of a total of 100) non-autonomous gene circuits fitted to data with non-zero diffusion terms leads to severe patterning defects (see S2 Figure for common patterning defects). This is likely due to numerical, not biological issues, since we do find circuit solutions that correctly reproduce gap gene patterns both in the presence and absence of diffusion using an alternative fitting approach: that fixes diffusion parameters *D^a^* to zero during optimization (see below). To further facilitate comparison with the static-Bcd model, we constrained the signs of regulatory weights to those reported in Manu *et al.* [23]. In previoius work, we have verified this network structure extensively against experimental data [18, 25, 26, 34]. Optimization was performed on the Mare Nostrum supercomputer at the Barcelona Supercomputing Centre (http://www.bsc.es). One optimization run took approximately 35 min on 64 cores.

The purpose of our reverse-engineering approach is not to sample parameter space systematically, but instead to discover whether there are specific model-fitting solutions that are consistent with the biological evidence and reproduce the dynamics of gap gene expression correctly. Global optimization algorithms are stochastic heuristics without guaranteed convergence, which means that for complex non-linear problems many optimization runs will fail or end up at sub-optimal solutions (see also discussions in [24, 26, 33]). In order to find the best-fitting solution, we therefore select solutions from 200 initial fitting runs as follows: (1) we discard numerically unstable circuits; (2) we only consider solutions with a root-mean-square (RMS) score less than 20.0 as most circuits with scores above this threshold show gross patterning defects; (3) we use visual inspection to detect remaining gross patterning defects among selected circuits (missing or bimodal domains, and disconnected boundaries. See S2 Figure) as previously described [34]. Out of the resulting 7 highest scoring circuits, only 3 recover the shifting dynamics of posterior gap domains. In order to rule out diffusion as a pattern-generating mechanism in these circuits, we compared their performance in the presence and absence of diffusion (see above). For this purpose, we used values of diffusion rates *D^a^* obtained by fitting our non-autonomous models with diffusion. All three circuits produce satisfactory gap gene patterns (including anteriorly shifting posterior trunk domains) whether diffusion is present or not. The best fit among these was selected for detailed analysis (see S1 Table, for parameter values).

The residual error of our best-fitting diffusion-less circuit (RMS = 10.73) lies at the lower end of the range of residual errors for fully-non-autonomous circuits with diffusion, which range from RMS scores of 10.43 to 13.32 [26]. This lends further support to the notion that diffusion is not essential for gap gene patterning. Moreover, our previous work also shows that circuits which were fit without weighting the data show somewhat lower RMS scores of 8.71 to 10.11 despite exhibiting more patterning defects at late stages [26]. The RMS score of the static-Bcd model (fit without weights) is higher, at 10.76 [22]. Taken together, this implies a slightly better quality-of-fit of our fully non-autonomous diffusion-less model compared to the static-Bcd diffusion-less circuits of Manu *et al.* [22].

### Gap gene circuit analysis

We characterise the time-variable geometry and topology of phase space in our fully non-autonomous gap gene circuit for every nucleus in a sub-range of the fitted model between 35 and 71% A–P position. This restricted spatial range allows us to simplify the analysis by excluding the influence of terminal gap genes *tll* and *hkb* on patterning (similar to the approach in [22]). We aim to identify those features of configuration space that govern the placement of domain boundaries, and thus the patterning capability of the gap gene system. We achieve this by generating instantaneous phase portraits for the model [43, 45] at 10 successive points in time (C13, C14A-T1–8, and gastrulation time). To generate an instantaneous phase portrait, all time-dependent parameter values—*i. e.* those corresponding to the profiles of external regulators—are frozen at every given time point. This yields an autonomous system for each point in time, for which we can calculate the position of steady states in phase space using the Newton-Raphson method [48, 49] as implemented by Manu *et al.* [23]. We classify steady states according to their stability, which is determined by the corresponding eigenvalues (see S1 Figure A).

Nuclei express a maximum of three trunk gap genes over developmental time, and only two at any given time point. Therefore, we project four-dimensional phase portraits into lower-dimensional representations to visualise them more easily. This yields a graphical time-series of instantaneous phase portraits for each nucleus, which allow us to track the movement, creation, and annihilation of steady states (typically attractors and saddles) by bifurcations. The transient geometry of phase space governs the non-autonomous trajectories of the system. We classify the dynamic behaviours exhibited by these trajectories into transitions, pursuits, and captures according to our previously established methodology (see Introduction and S1 Figure B) [43].

## Results

### Non-autonomous gap gene circuits without diffusion

Previously published non-autonomous gap gene circuits suggest a specific regulatory structure for the gap gene network in *D. melanogaster* (Fig. 3A) [26]. This structure is consistent with the network predicted by the static-Bcd model of Manu *et al.* [23], and with the extensive genetic and molecular evidence available in the published literature on gap gene regulation [18]. Unfortunately, it is difficult to derive insights about dynamic regulatory mechanisms from a static network diagram. Computer simulations help us understand which network interactions are involved in positioning specific expression domain boundaries across space and time [24–26, 34]. Although powerful, this simulation-based approach has its limitations. It cannot tell us how expression dynamics are brought about: for instance, why some gap domain boundaries remain stationary while others shift position over time. To gain a deeper understanding of the underlying regulatory dynamics, we analyse the configuration space of a fully non-autonomous gene circuit through instantaneous phase portraits (S1 Figure B) [43], analogous to the autonomous phase-space analysis presented by Manu and colleagues [23] (Fig. 2). This type of analysis requires diffusion-less gap gene circuits to keep the dimensionality of phase space at a manageable level.

**Figure 3.**
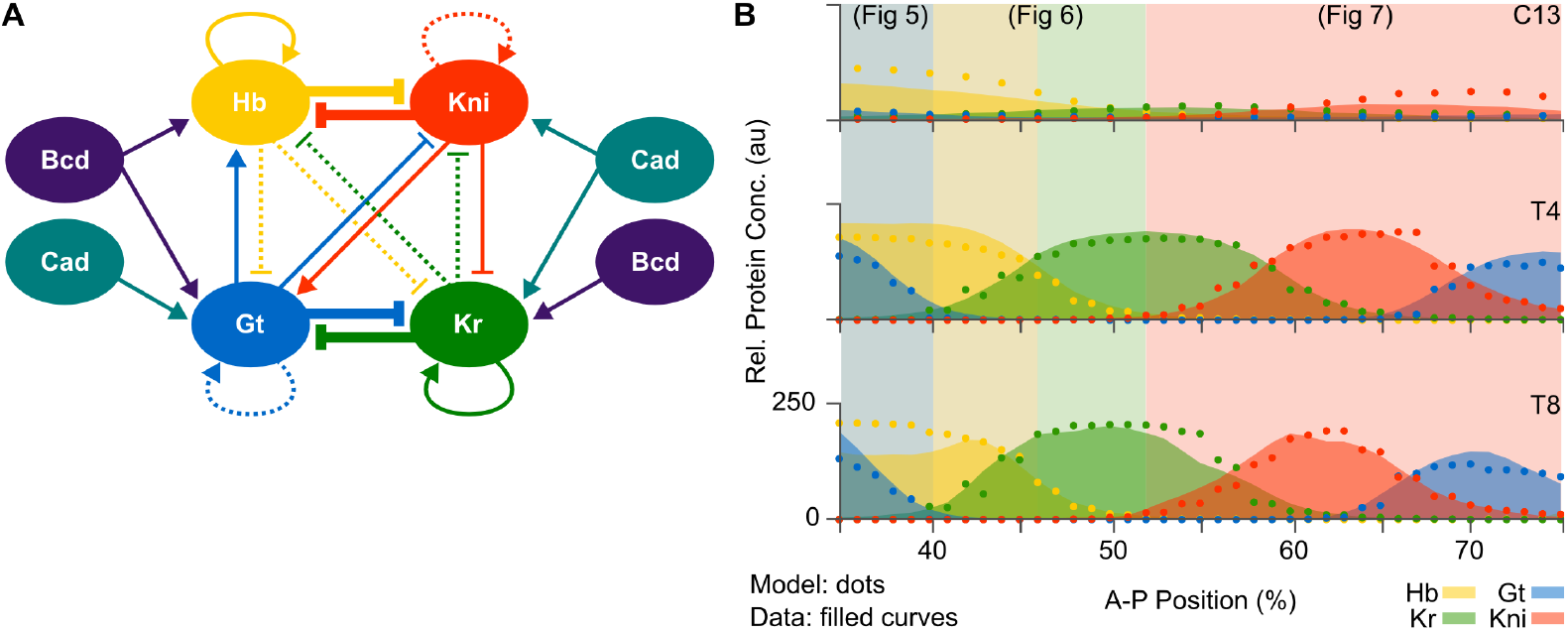
Non-autonomous gap gene circuits. **(A)** Regulatory structure derived from previously published non-autonomous gap gene circuits with diffusion [26]. Connecting arrows and T-bars represent activating and repressive interactions, respectively. Line thickness indicates an interaction's relative strength, with very weak interactions dashed. **(B)** Model output of a fully non-autonomous gene circuit without diffusion (dots) and gap protein data (filled curves) at three time points: C13 (early), T4 (mid), and T8 (late blastoderm stage). The x-axis represents %A–P position, where 0% is the anterior pole. The y-axis represents relative protein concentration in arbitrary units (au). Coloured background areas indicate different dynamic patterning mechanisms that are shown in detail in Figs. 5–7.

We obtained fully non-autonomous gap gene circuits that lack diffusion through model fitting with diffusion parameters *D^a^* fixed to zero and interaction signs constrained to those of previous works (as described in “Models and Methods”). This resulted in a set of three selected, well-fitted circuits. The network topology of these gene circuit models correspond to that shown in Fig. 3A. The following analysis is based on the best-fitting model with a root mean square (RMS) residual error of 10.73, which constitutes a slight overall improvement in quality-of-fit compared to static-Bcd models (see “Models and Methods” and [22, 26]). Its regulatory parameter values are listed in S1 Table.

This diffusion-less non-autonomous gene circuit accurately reproduces gap gene expression (Fig. 3B). In particular, it exhibits correct timing and relative positioning of domain boundaries. Together with the fact that it fits the data equally well as equivalent circuits with diffusion (see “Models and Methods”, and [26]), this confirms earlier indications that gap gene product diffusion is not essential for pattern formation by the gap gene system [23, 25]. Interestingly, previously published diffusion-less static-Bcd circuits show rugged patterns with abrupt “on/off” transitions in expression levels between neighbouring nuclei [23]. In contrast, diffusion-less fully non-autonomous circuits produce smooth spatial expression patterns with a graded increase or decrease in concentration levels across domain boundaries. This is because non-autonomy, with its associated movement of attractors and separatrices over time, provides increased flexibility for fine-tuning expression dynamics over time compared to models with constant phase-space geometry (see below). In biological terms, it suggests that the expression of smooth domain boundaries does not strictly require diffusion. Although diffusion undoubtedly contributes to this process in the embryo, its role may be less prominent than previously thought [23,25].

### Maternal gradient decay affects the level and timing of gap gene expression

We used our non-autonomous gap gene circuit to assess the effect of maternal gradient decay on gap gene regulation. One way to isolate this effect is to compare the output of the fully non-autonomous model—with decaying maternal gradients—to simulations using the same model parameters, but keeping maternal gradients fixed to their concentration levels early during the blastoderm stage (time class C12). As shown in Fig.4, the relative order and positioning of gap domains remain unaffected when comparing models with fixed versus time-variable gradient concentrations. This indicates that maternal gradient decay is not strictly required for correct pattern formation by gap genes.

**Figure 4.**
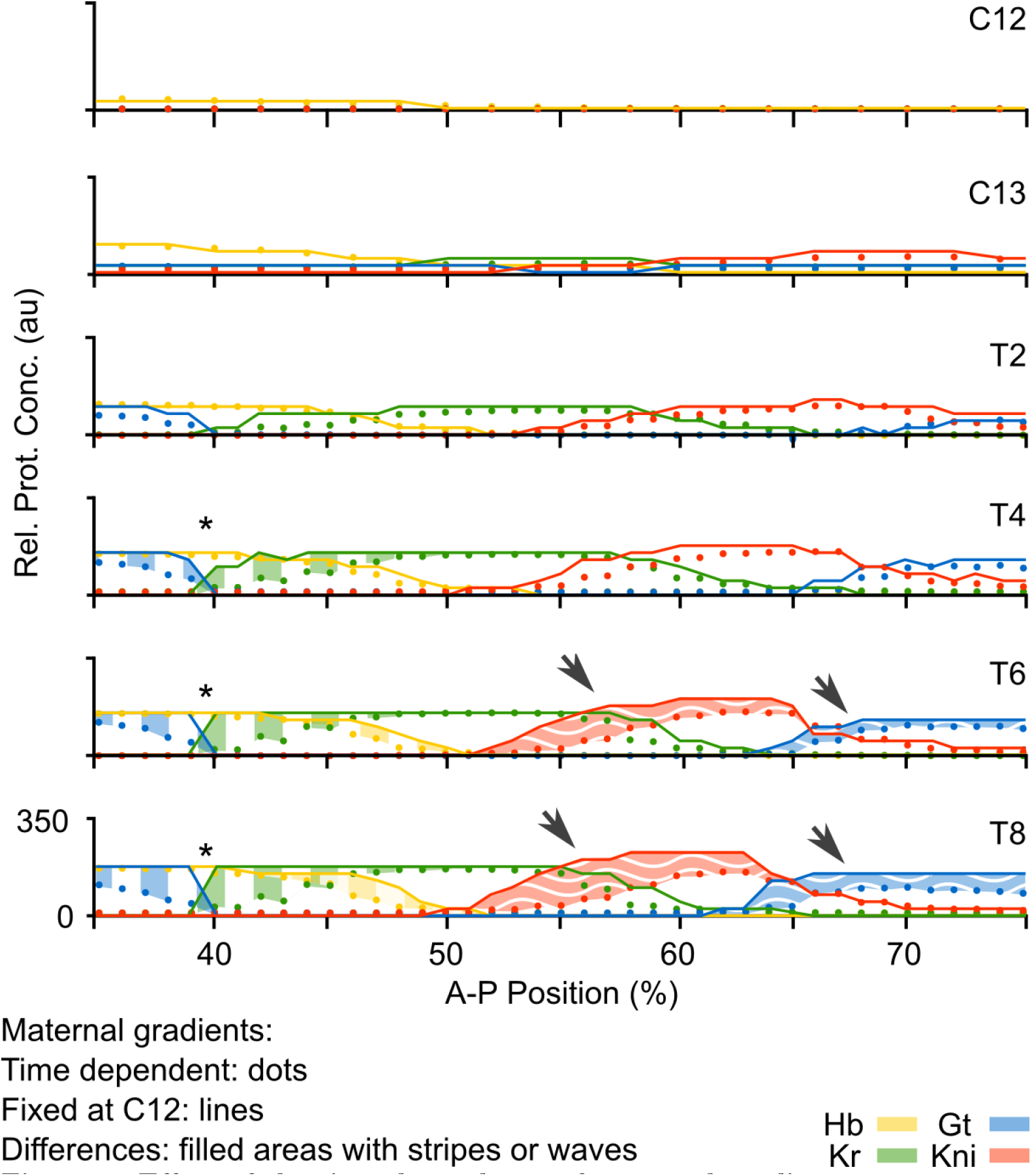
Effect of the time-dependence of maternal gradients on gap gene pattern formation. **(A)** Plots show output from the non-autonomous gap gene circuit with time-variable maternal gradients (dots), compared to output from the same model with maternal gradients fixed to their values at cycle C12 (early blastoderm stage; lines). Y-axes represent relative protein concentrations in arbitrary units (au); x-axes represents %A–P position, where 0% is the anterior pole. Differences between the two model simulations are shaded using vertical stripes in the anterior trunk region, and wavy horizontal stripes in the posterior. Asterisks mark over-expression in the region of the Gt/Kr interface; arrows mark “overshoot” of gap domain shifts in the posterior of the embryo.

We do observe, however, that maternal gradient dynamics significantly affect the levels of gap gene expression throughout the trunk region of the embryo (Fig. 4, shaded areas). While early expression dynamics are very similar in both models (time classes C12–T2), they begin to diverge at later stages. The fully non-autonomous model reaches peak expression at T2/T4, but the autonomous model without maternal gradient decay overshoots observed expression levels in the data between T4 and T8. This indicates that maternal gradient decay leads to decreasing activation rates at the late blastoderm stage, thereby regulating the timing and level of peak gap gene expression. Such a limiting regulatory effect of maternal gradients has been proposed before [25, 42], but has never been tested explicitly.

Interestingly, the overshoot occurs in different ways in the anterior and the posterior of the embryo. In the anterior, maximum concentrations of Hb and Kr across each domain remain unchanged, but levels of expression keep increasing around the Kr/Gt interface, rendering the domain boundaries steeper and less smooth in the simulation without maternal gradient decay (Fig. 4, asterisk). In the posterior, we observe increased levels of Kni and Gt across large parts of their respective expression domains (Fig. 4, arrows). These effects are asymmetric: both posterior Kni and Gt domains exhibit an anterior expansion, while the posterior boundary of the Kni domain is not affected. Considering that both of these domains shift towards the anterior over time (Fig. 1) [25, 42], we interpret this as follows: maternal gradient decay not only decreases the rate of expression at late stages in the posterior region, but also leads to a slow-down of gap domain shifts, thereby limiting the extent of the shift. In the autonomous simulation without maternal gradient decay, both Kni and Gt domains keep on moving, which explains the observed expansion and increase of expression levels towards the anterior part of the domain.

### Non-autonomous regulatory mechanisms for gap gene patterning

We asked whether the differing effects of maternal gradient decay in the anterior and the posterior of the embryo depend on the presence of different regulatory mechanisms in these regions [23]. To validate this hypothesis, we need to understand and characterise the dynamic mechanisms underlying gene regulation in our non-autonomous model. We achieve this through analysis of the time-variable phase spaces of nuclei across the trunk region of the embryo using the methodological framework presented in the Introduction (S1 Figure B; see [43] for details). To briefly reiterate, this analysis is based on the characterization of the changing phase space geometry that shapes the trajectories of the system. The shape of a trajectory indicates typical dynamical behaviors, that can be classified into four distinct categories—transitions, pursuits, as well as geometrical and topological captures—each showing particular dynamic characteristics. These categories provide mechanistic explanations for the dynamic behavior of the system. For every nucleus, we then compare these non-autonomous mechanisms to the autonomous mechanisms of pattern formation found in the static-Bcd model [23]. This direct comparison allows us to identify the causes underlying the observed effects of maternal gradient decay on the temporal dynamics of gap gene expression.

In agreement with Manu *et al.* [23], we find different patterning modes anterior and posterior to 52% A–P position. Just like in static-Bcd models, anterior expression dynamics are governed by convergence of the system towards attractors in a multi-stable regime. In contrast, our model differs from that of Manu *et al.* [23] concerning posterior gap gene regulation. We find that a monostable spiral sink drives gap domain shifts in the posterior of the embryo; this differs markedly from the unstable manifold observed in static-Bcd gap gene circuits [23]. An in-depth analysis and biological discussion of spatial pattern formation driven by this mechanism goes beyond the scope of this study. It is provided elsewhere [44]. Here, we focus on temporal aspects of gene regulation and pattern formation, namely the regulation of the velocity of gap domain shifts by maternal gradient dynamics in the posterior of the embryo.

#### Anterior non-autonomous mechanisms of pattern formation

Phase portraits of nuclei in the anterior of the embryo (35 to 51% A–P position) are multi-stable at every time point. Every instantaneous phase portrait contains multiple attractors. Distinct attractors govern the dynamics of gap gene expression at different points in space and time. We identify three alternative non-autonomous mechanisms which control the positioning of domain boundaries in the anterior trunk region of the embryo.

The posterior border of the anterior Gt domain forms between 35 and 40% A–P position (Fig. 5A). Of all the trunk gap genes, nuclei in this region of the embryo only express *hb* and *gt*. Gap gene expression dynamics are governed by the same attractor across different nuclei (Fig. 5B). Each trajectory starts at non-zero (maternal) Hb concentration and initially converges towards the attractor located at high Hb and Gt concentrations (Fig. 5B). The phase portrait for every nucleus changes in the non-autonomous simulation as maternal gradients decay. For the nuclei between 35 and 40% A–P position, the attractor drops towards lower Gt levels over time, while maintaining high concentrations of Hb. Convergence towards the moving attractor is shaping these trajectories (Fig. 5B, grey trajectories). At some point, the attractor “overtakes” (*i. e.* passes in front of) the trajectory in phase space, which leads to a marked change in the trajectory's direction. Although all nuclei across the Gt boundary show qualitatively similar behaviour, the timing of attractor movement differs markedly from one nucleus to another. The further posterior a nucleus is located along the A–P axis, the earlier the drop of the attractor occurs (Fig. 5B). As a result, non-autonomous trajectories bend towards low Gt levels at increasingly early stages as we move towards the posterior, which results in lower overall Gt concentration profiles as we proceed from 35 to 39% A–P position. This causes a gradual decrease in Gt concentration along the Gt boundary in the non-autonomous model which results in a smooth boundary, even in the absence of diffusion. In the phase portraits of nuclei at 37 and 39%, we observe a saddle-node bifurcation (at T7 and T8, respectively) which annihilates the attractor to which the trajectory is initially converging. However, this bifurcation occurs too late to perceivably affect the dynamics of the system. We conclude that the position of the posterior boundary of the anterior Gt domain is largely defined by the timing of attractor movement. Therefore, it is governed by what we call a pursuit mechanism in S1 Figure B [43].

**Figure 5.**
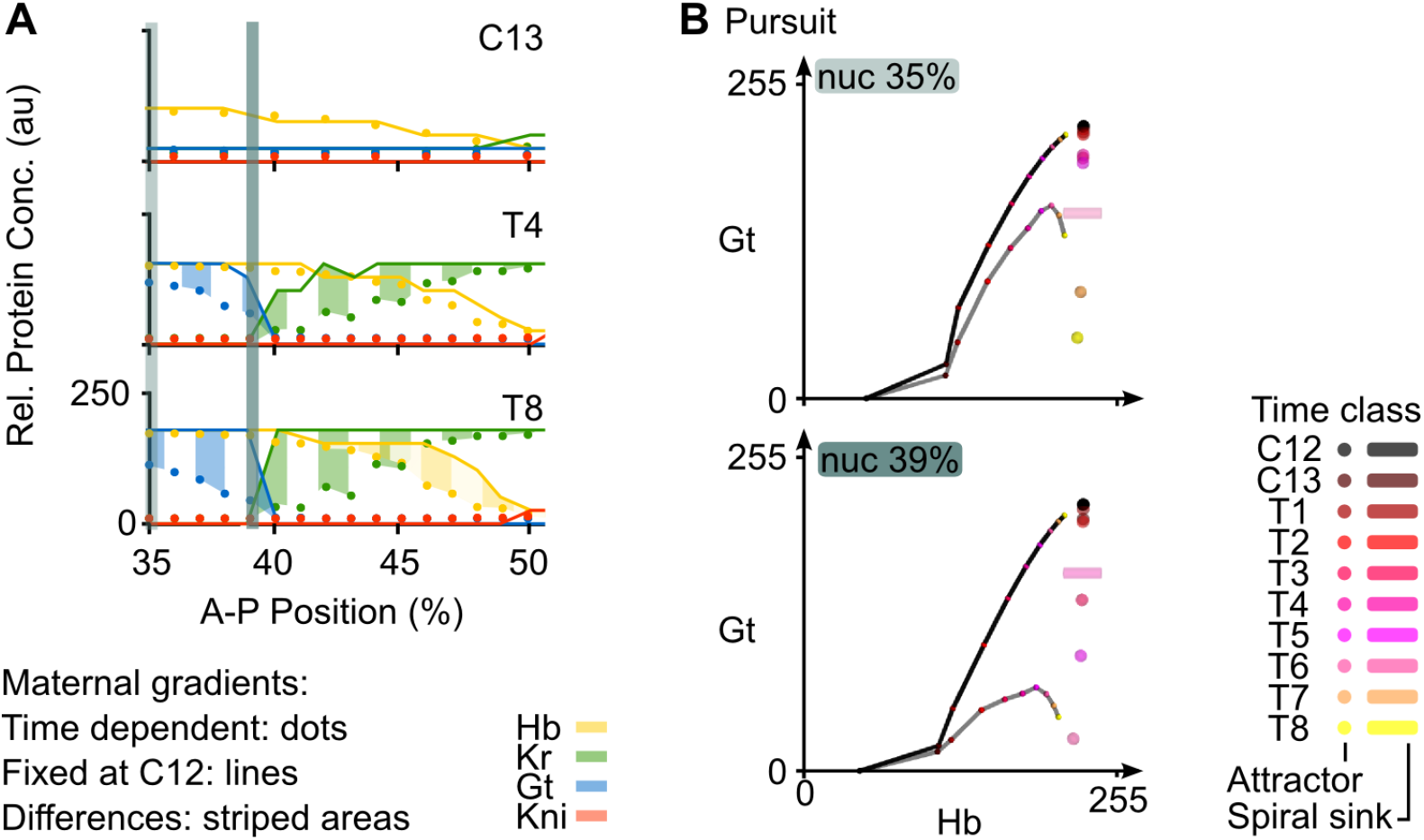
Positioning the posterior boundary of the anterior Gt domain. **(A)** Output of the non-autonomous gene circuit (dots) versus the same model without maternal gradient decay (lines) shown at cleavage cycle C13 and C14A (time classes T4 and T8) for nuclei within 35–52% A–P position. Axes and colouring scheme as in Fig. 3B. Blue vertical bars mark the nuclei at 35% and 39%A–P position shown in (B). **(B)** Phase portraits for nuclei at 35% (top) and 39 %A–P position (bottom). Phase portraits are shown as two-dimensional projections onto the plane defined by Hb (x-axis) and Gt (y-axis) concentrations (in arbitrary units, au). Non-autonomous trajectories shown as grey lines and autonomous trajectories as black lines. Attractors shown as spheres (point attractors) and cylinders (indicating a spiral sink). Small coloured dots on trajectories indicate the position in space of that trajectory at different time points. Colouring of attractors and trajectory positions indicates time class (see key). Other steady states have been omitted for clarity, since they do not shape the trajectories in these nuclei. See text for details.

In contrast, the simulation without gradient decay does not show a drop in attractor position, since the phase portrait does not change over time and the attractor remains at high Hb and Gt concentration until the onset of gastrulation (Fig. 5B, black trajectory and black steady state). This causes its trajectory to increasingly diverge from the non-autonomous case, explaining the elevated Gt concentrations in this region of the embryo (Fig. 4).

Further posterior, in the region between 40 and 52% A–P position, the only gap genes that are expressed are *hb* and *Kr*. In this area, the posterior boundary of the anterior Hb domain and the anterior boundary of the central Kr domain overlap (Fig.6A). In the non-autonomous model, this boundary interface is set up by two different regulatory mechanisms (Fig. 6B,C). Phase portraits of nuclei between 41 and 45% A–P position (Fig. 6A, yellow vertical bar) show the following dynamics: for most of the time, system trajectories converge towards an attractor located at high Hb and high Kr concentration (Fig. 6B, grey trajectory). However, these trajectories are transient and remain at low Kr and intermediate Hb concentrations, far from steady state. At T7, two simultaneous saddle-node bifurcations give rise to two new attractors, one at high Hb and the other at high Kr concentration (Fig. 6B). Two new saddles are also created. System trajectories are caught in the basin of the attractor with high Hb levels. This only has a noticeable effect in more posterior nuclei (*e. g.* at 43% A–P position in Fig. 6B), where there is a drastic (but late) change in the direction of the trajectory. At T8, two additional saddle-node bifurcations occur, which annihilate the high Hb/high Kr attractor, as well as the newly created attractor at high Hb. This leaves only the attractor at high Kr concentrations (Fig. 6B). Trajectories of the system are once again caught in a different basin of attraction. However, this second round of bifurcations occurs too late to still have a substantial effect on expression dynamics. A non-autonomous trajectory being caught in a new basin of attraction due to a preceding bifurcation event is called a topological capture (S1 Figure B) [43].

**Figure 6.**
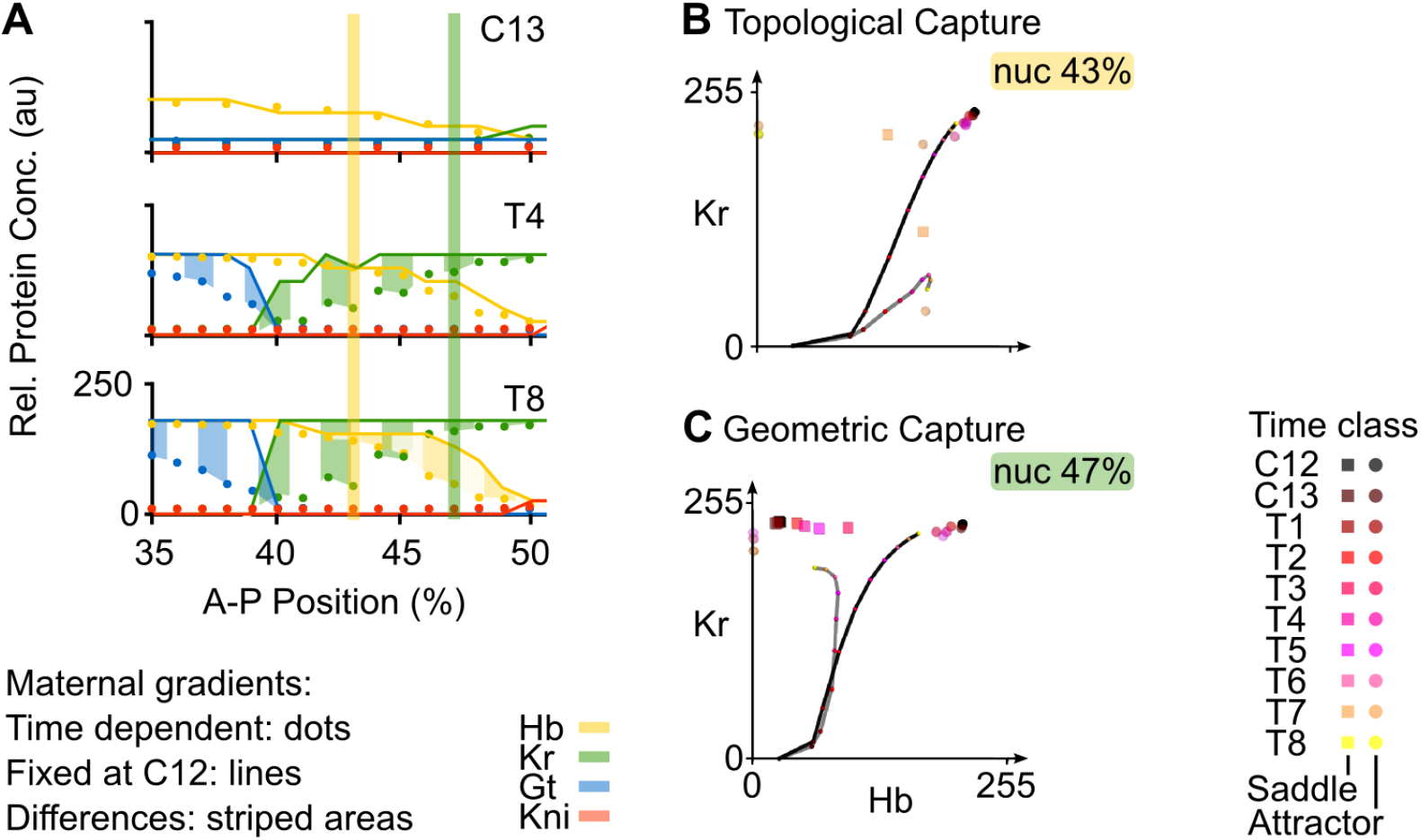
Positioning the Hb-Kr interface. Output of the non-autonomous gene circuit (dots) versus the same model without maternal gradient decay (lines) shown at cleavage cycle C13 and C14A (time classes T4 and T8) for nuclei between 35–52% A–P position. Axes and colouring scheme as in Fig. 3B. Yellow and green vertical bars mark the nuclei at 43% and 47%A–P position shown in (B) and (C) respectively. **(B)** Phase portrait for nucleus at 43% position. **(C)** Phase portrait for nucleus at 47% position. Phase portraits are shown as two-dimensional projections onto the plane defined by Hb (x-axis) and Kr (y-axis) concentrations (in arbitrary units, au). Non-autonomous trajectories are shown as grey lines and autonomous trajectories, as black lines. Point attractors are represented by spheres and saddles points by squares. Small coloured dots on the trajectories indicate the position in phase space (Hb and Kr concentrations) of the trajectory at different time points. Colouring of attractors and trajectory positions indicates time class (see key). Other steady states have been omitted for clarity, since they do not shape trajectories in these nuclei. See text for details.

In the region between 46 and 52% A–P position (Fig. 6A, green vertical bar), we observe a different kind of dynamical behaviour. Similar to more anterior nuclei, these instantaneous phase portraits have an attractor at high Hb and high Kr levels and trajectories converge to this steady state at early stages (Fig. 6C). In contrast to more anterior nuclei, however, there is a saddle located on the Hb-Kr plane. Between time class T2 and T6, the position of this saddle moves towards higher Hb levels. When a saddle moves on a phase portrait, it drags the associated separatrix with it (S1 Figure A). These concerted movements change the location of the boundaries between existing basins of attraction. When a separatrix “overtakes” a trajectory in phase space, a geometrical capture occurs (S1 Figure B) [43]. This can be observed in the nucleus at 47% A–P position (Fig. 6C, grey trajectory). Here, the trajectory gets captured by the moving separatrix between T2 and T5, and later starts to converge towards the attractor at high Kr, limiting Hb concentrations at intermediate levels. Taken together, our results indicate that the posterior boundary of the anterior Hb domain, as well as the anterior boundary of the central Kr domain, are positioned by a combination of topological and geometrical capture events.

In simulations without gradient decay, captures cannot occur (Fig. 6B and C, black trajectory and black steady state). In both nuclei at 43 and 47%, trajectories keep on converging towards the attractor at high Hb and Kr. This results in higher and sustained Hb and Kr levels throughout the region where the two factors are co-expressed. It explains why there are very abrupt boundaries between Gt and Kr, as well as between Hb and Kni, instead of the smooth interfaces between the corresponding domains observed in the non-autonomous model (Fig. 6A).

Taken together, our evidence suggests that the non-autonomous mechanisms positioning anterior gap domains are equivalent to the corresponding autonomous mechanisms from the static-Bcd model described by Manu *et al.* [23] since they too rely on attractor position and/or switching between basins of attraction. In their work, just as in ours, the Gt boundary is set by an attractor moving from high to low Gt concentrations (across space, *i.e.* moving along the A–P axis), and the Hb/Kr interface is positioned by attractor selection: nuclei anterior to this border fall into the basin of an attractor with high Hb, nuclei posterior of the border end up in the basin of an attractor with high Kr concentration. Instead of a static switch, however, we find nuclei being captured by different basins at different time points across space. Still, the overall principle of boundary placement by attractor selection remains the same between static-Bcd and fully non-autonomous gap gene circuit models. The fact that similar regulatory principles are at work in both models validates our approach, and confirms that the placement of stationary domain boundaries in the anterior of the embryo does not depend in any fundamental way on the dynamics of maternal inputs.

#### Posterior non-autonomous mechanisms of pattern formation

Expression boundaries posterior to 52% A–P position are not stationary but move towards the anterior over time, causing a shift and concurrent narrowing of gap domains in this region (Fig. 7A) [25, 42]. Surprisingly, we find that these shifting posterior gap domains are governed by quite different phase space geometries in our model compared to those previously reported. Manu *et al.* [23] found that posterior gap gene expression dynamics are controlled by an unstable manifold embedded in a multi-stable phase space geometry in their static-Bcd model. In contrast, our fully non-autonomous gap gene circuit features no such manifold: the phase portraits of posterior nuclei lack saddle points since they are monostable throughout the blastoderm stage and only contain a single attractor (Fig. 7B). This attractor is not a regular point attractor. Its complex eigenvalues reveal that it is a *spiral sink* (also known as a *focus*; see S1 Figure A) ( [36]). Like regular point attractors, spiral sinks are stable, in that they draw trajectories asymptotically towards them. Unlike regular point attractors, these trajectories do not approach the steady state in a straight line, but rather spiral inward towards the sink. Since sinks are a consistent feature of the phase spaces of nuclei in the posterior of the embryo, it is likely that they are important for the spiral-shaped geometry of the trajectories observed in this region (Fig. 7B). The spiral geometry in turn is responsible for the ordered succession of transient gap gene expression governing dynamic domain shifts. A full characterization of this patterning mechanism is presented elsewhere [44]. For the purpose of our present analysis, we conclude that the non-autonomous mechanism patterning the posterior region corresponds to a pursuit, where the system follows but never reaches a moving attractor (S1 Figure B).

**Figure 7.**
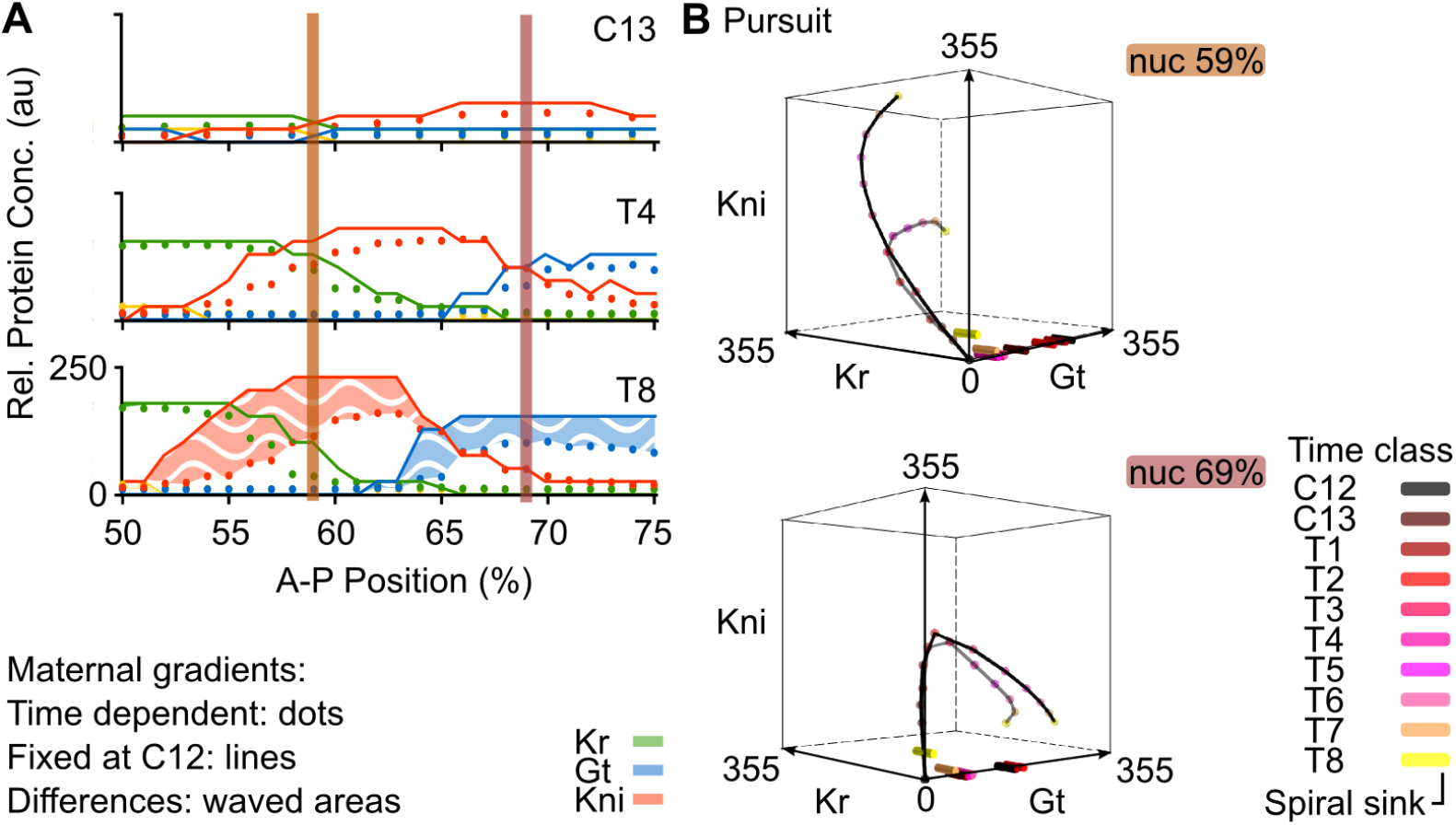
Regulating the extent and timing of posterior gap domain shifts. **(A)** Output of the non-autonomous gene circuit (dots) versus the same model without maternal gradient decay (lines) shown at cleavage cycle C13 and C14A (time classes T4 and T8) for nuclei between 50–75% A–P position. Axes and colouring scheme as in Fig. 3B. Red vertical bars mark the nuclei at 59% and 69%A–P position shown in (B). **(B)** Phase portraits for nuclei at 59% (top) and 69 %A–P position (bottom). Phase portraits are shown as three-dimensional projections onto the sub-space defined by Kr (x-axis), Gt (y-axis) and Kni (z-axis) concentrations (in arbitrary units, au). Non-autonomous trajectories shown as grey lines and autonomous trajectories as black lines. Spiral sinks are represented by cylinders. Small coloured dots on trajectories indicate the position in phase space of the trajectory at different time points. Colouring of attractors and trajectory positions indicates time class (see key). See text for details.

The correct geometry of transient trajectories in the posterior of the embryo depends crucially on maternal gradient decay. As we can see in Fig. 7B (black trajectories), simulations without dynamic gradient concentrations show much less tightly wound spirals. This means that the transition between the expression of successive gap genes in this region is delayed. For example, the nucleus at 59% A–P position shows a delayed down-regulation of *Kr*, while *kni* keeps on accumulating. This provides a straightforward explanation of the “overshoot” of Kni and Gt domain shifts observed in the simulation without maternal gradient decay (Fig. 7A). In biological terms, it suggests that the disappearance of Cad from the abdominal region of the embryo is required for correct pattern formation, by limiting the timing—and as a result, the extent—of gap domain shifts.

## Discussion

In this paper, we have examined the explicitly time-dependent aspects of morphogen gradient interpretation by a gene regulatory network; the gap gene system of the vinegar fly *D. melanogaster*. Using a fully non-autonomous gap gene circuit, we compared the dynamics of gene expression in the presence and absence of maternal gradient decay. We find that dynamic changes in the concentration of maternal morphogens Bcd and Cad affect the timing and rate of gap gene expression. The precise nature of these effects differs between the anterior and the posterior region of the embryo. In the anterior, gradient decay creates smooth domain borders by preventing the excessive accumulation of gene products across boundary interfaces between neighbouring gap domains. In the posterior, gradient decay limits the rate of gap gene expression, and therefore the extent of gap domain shifts, towards the end of the blastoderm stage. A temporal effect on gene expression rates is translated into slowing rates of domain shifts, which in turn alter the spatial positioning of expression boundaries. As a consequence, gradient decay stabilises spatial gap gene patterns before the onset of gastrulation. An effect of maternal gradient decay on gap gene expression rates has been suggested before—based on the analysis of quantitative expression data [25, 42]. However, only mechanistic dynamical models—such as the non-autonomous gap gene circuits presented here—can provide specific mechanisms and quantitative causal evidence for this aspect of gap gene regulation.

Our analysis suggests that maternal gradient decay—specifically, the disappearance of Cad from the abdominal region of the embryo—has an important role in regulating the timing of gap gene expression as well as limiting the rate and extent of gap domain shifts in the posterior of the embryo. This result is consistent with experimental data indicating that Cad affects gap domain shifts. Mutants lacking maternal *cad*, which show a reduced level of Cad protein throughout the blastoderm stage [28], show a delay in the shift of the posterior domains of *kni* and *gt* [32,44]. However, Cad does not seem to act exclusively. An indirect role of Bcd in regulating gap domain shifts through altering gap-gap interactions was suggested by a modelling study [30]. It remains unclear whether Cad is also involved in mediating this effect. Finally, a recent study of Bcd-dependent regulation of *hb* postulated an additional mechanism for gap gene down-regulation that acts before maternal gradient decay occurs [2]. This could have an indirect effect on the timing of late (Bcd-independent) *hb* regulation, which may mediate the direct effect of Bcd decay on late *hb* expression we are observing in our models.

To better understand the mechanistic basis for the observed differences in patterning between the anterior and the posterior, we analysed the time-variable phase portraits in our non-autonomous model [43]. In agreement with a previous study based on autonomous phase space analysis of static-Bcd gap gene circuits [23], we find that two distinct dynamical regimes govern gap gene expression anterior and posterior to 52% A–P position (Fig. 8). Stationary domain boundaries in the anterior are governed by regulatory mechanisms that are equivalent in static-Bcd and fully non-autonomous models (our work and [23]): they take place in a multi-stable dynamical regime where the posterior boundary of the anterior Gt domain is set by the movement of an attractor in phase space, and the posterior boundary of the anterior Hb domain is set by attractor selection (*i. e.* the capture of transient trajectories in the non-autonomous case) (Fig. 8, left). Attractor movement in fully non-autonomous models leads to smooth expression boundaries, which are absent in the static-Bcd case. In contrast, static-Bcd and non-autonomous models suggest different mechanisms for gap domain shifts in the posterior of the embryo. While these shifts are controlled by an unstable manifold in the static-Bcd gene circuit model [23], we find a pursuit mechanism featuring a monostable spiral sink to govern their behaviour in our fully non-autonomous analysis (Fig. 8). The spiralling geometry of transient trajectories imposes temporal order on the progression of gap genes being expressed. If arranged appropriately across nuclei in the posterior of the embryo, this temporal progression from *Kr* to *kni* to *gt* to *hb* leads to the emergence of the observed kinematic domain shifts [44].

**Figure 8.**
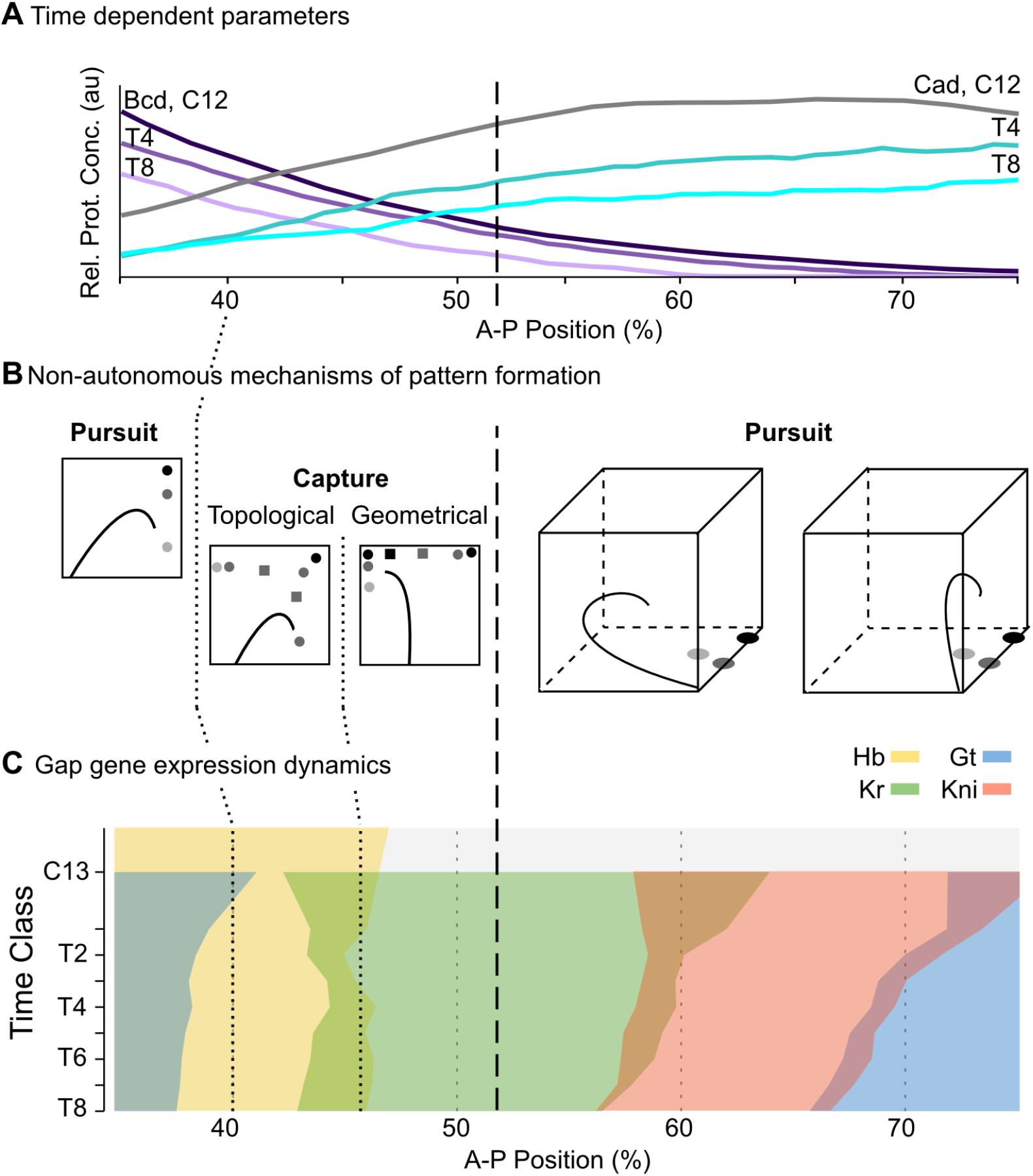
Summary of non-autonomous mechanisms for gap gene pattern formation in *D. melanogaster*. **(A)** Non-autonomous gap gene circuits implement realistic, time-dependent dynamics of maternal morphogen gradients (Bcd in purple, Cad in cyan). Y-axis shows relative protein concentration (in arbitrary units, au); X-axis shows %A–P position, where 0% is the anterior pole. **(B)** Different non-autonomous mechanisms of pattern formation are active at different positions along the A–P axis of the embryo. Stylized projections of phase space are shown. See Box.1B and [43] for nomenclature. **(C)** Gap gene expression dynamics differ between the anterior and the posterior regions of the embryo. While domain boundaries in the anterior are stationary, boundaries in the posterior shift towards the anterior over time. Time-space plot as in Fig.1A: note that time flows downward along the y-axis (cycle C13 and time classes T1-8 as defined in Models and Methods). The dashed vertical line spanning all panels indicates a bifurcation event at 52% A–P position, which separates the multi-stable from the oscillatory regime.

It is important to note that similar regulatory principles can be found in all three solutions of our fully non-autonomous model that reproduce gap-gene patterning correctly both in the presence and absence of diffusion. We have chosen the most structurally stable solution for detailed analysis. The other two circuits show more variability of regulatory features both across space and time. Still, both of these models consistently exhibit multi-stability in the anterior, and spiral sinks as well as transiently appearing and disappearing limit cycles in the region posterior to 52% A–P position. This indicates that the two main dynamical regimes described here—stationary boundaries through attractor selection in the anterior vs. shifting gap domain boundaries through spiralling trajectories in the posterior—are reproducible across model solutions.

It is important to note that non-autonomy of the model is not strictly required for the spiral sink mechanism to pattern the posterior of the embryo. Simulations with fixed maternal gradients demonstrate that domain shifts can occur in an autonomous version of our gap gene circuit (see Figs. 4 and 7). The reason why earlier models [22, 23] do not feature spiral sinks remains unknown although one possibility is that fitting in the absence of diffusion somehow benefits characterisations of posterior pattern formation in terms of oscillatory behaviours. In spite of this, there are two reasons to consider the mechanism proposed here an important advance over the unstable manifold proposed by Manu *et al.* [23]. The first reason is technical: non-autonomous gap gene circuits—implementing correct maternal gradient dynamics—are more accurate and stay closer to the data than the previous static-Bcd model. The fact that the quality of a reverse-engineered model usually depends on the quality of its fit to data implies that our model provides more accurate and rigorous predictions than previous efforts. The second reason is conceptual: although it is difficult to interpret an unstable manifold in an intuitive way, it is straightforward to understand the spiral sink as a damped oscillator patterning the posterior of the embryo. The presence of an oscillatory mechanism in a long-germband insect such as *D. melanogaster* has important functional and evolutionary implications, which are discussed elsewhere [44].

Analysis of an accurate, non-autonomous model is required to isolate and study the explicitly time-dependent aspects of morphogen interpretation by the gap gene system. Here, we have shown that such an analysis is feasible and leads to relevant and specific new insights into gene regulation. Other modelling-based studies have used non-autonomous models before (see, for example, [16, 26, 34, 50–53]). However, none of them have directly addressed the proposed role of non-autonomy in pattern formation [17]. Our analysis provides a first step towards a more general effort to transcend this limitation in our current understanding of the dynamic regulatory mechanisms underlying pattern formation during animal development.

## Supporting Information

**Table S1.**
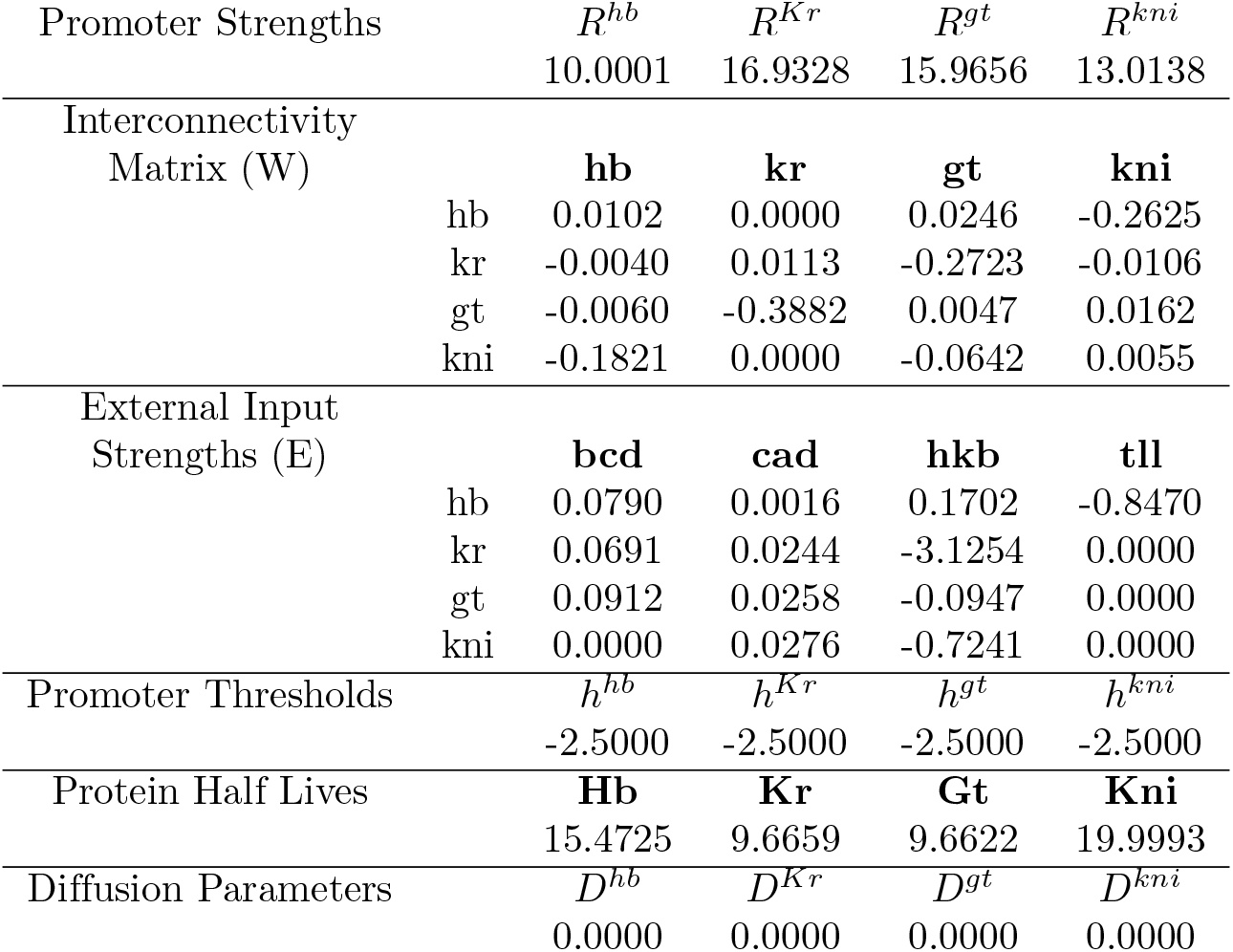
**Values of the parameters in the non-autonomous gap gene circuit model** Model equations are shown in the Models and Methods section.

|  |  |  |  |  |  |
| --- | --- | --- | --- | --- | --- |
| Promoter Strengths | | $R^{hb}$ | $R^{Kr}$ | $R^{gt}$ | $R^{kni}$ |
|  |  | 10.0001 | 16.9328 | 15.9656 | 13.0138 |
| Interconnectivity Matrix (W) |  | **hb** | **kr** | **gt** | **kni** |
|  | hb | 0.0102 | 0.0000 | 0.0246 | -0.2625 |
|  | kr | -0.0040 | 0.0113 | -0.2723 | -0.0106 |
|  | gt | -0.0060 | -0.3882 | 0.0047 | 0.0162 |
|  | kni | -0.1821 | 0.0000 | -0.0642 | 0.0055 |
| External Input Strengths (E) |  | **bcd** | **cad** | **hkb** | **tll** |
|  | hb | 0.0790 | 0.0016 | 0.1702 | -0.8470 |
|  | kr | 0.0691 | 0.0244 | -3.1254 | 0.0000 |
|  | gt | 0.0912 | 0.0258 | -0.0947 | 0.0000 |
|  | kni | 0.0000 | 0.0276 | -0.7241 | 0.0000 |
| Promoter Thresholds | | $h^{hb}$ | $h^{Kr}$ | $h^{gt}$ | $h^{kni}$ |
|  |  | -2.5000 | -2.5000 | -2.5000 | -2.5000 |
| Protein Half Lives |  | **Hb** | **Kr** | **Gt** | **Kni** |
|  |  | 15.4725 | 9.6659 | 9.6622 | 19.9993 |
| Diffusion Parameters | | $D^{hb}$ | $D^{Kr}$ | $D^{gt}$ | $D^{kni}$ |
|  |  | 0.0000 | 0.0000 | 0.0000 | 0.0000 |

**Figure S1.**
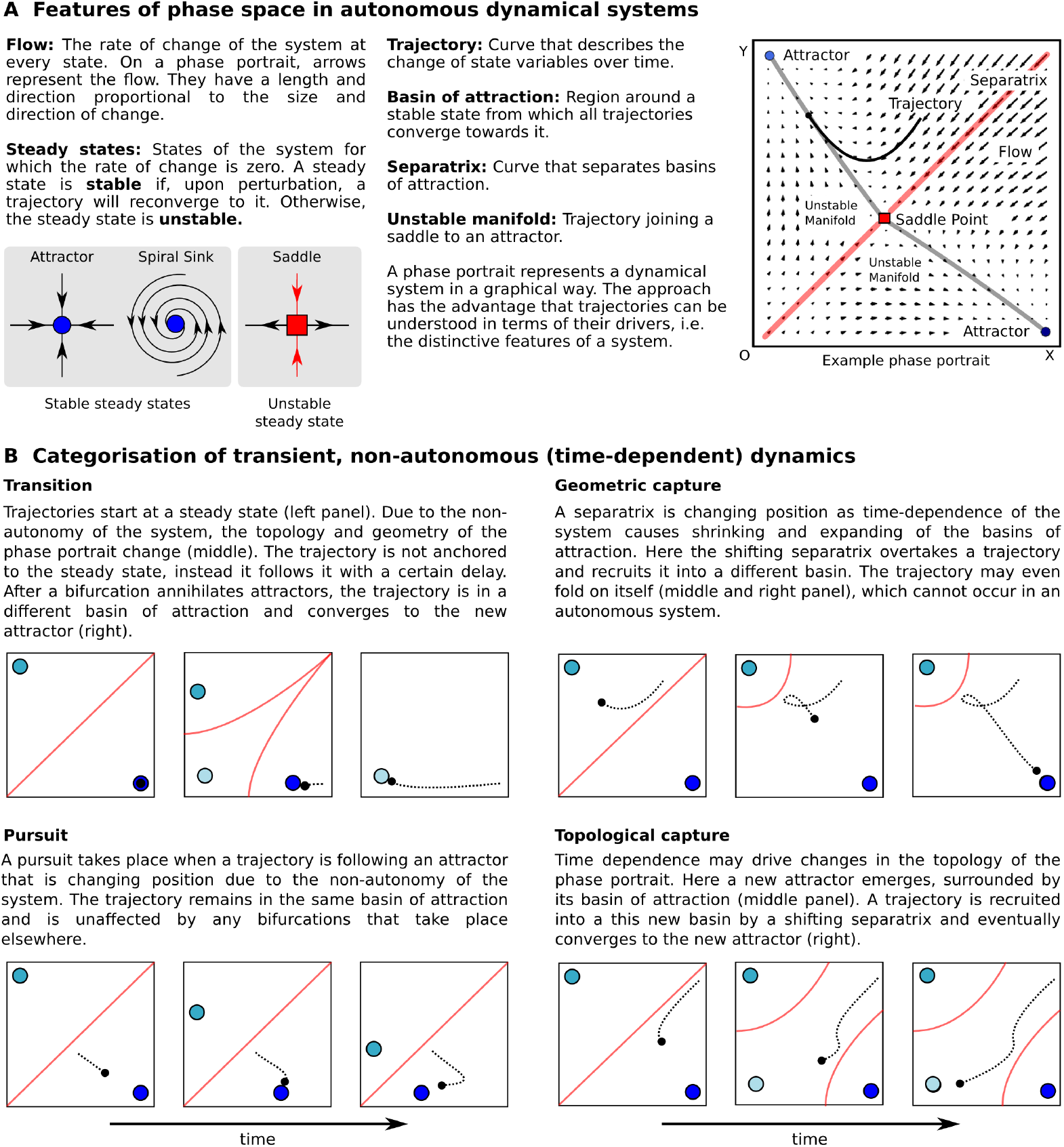
Dynamical systems concepts. **(A)** Features of phase space in autonomous dynamical systems. **(B)** Categorisation of transient, non-autonomous dynamics.

**Figure S2.**
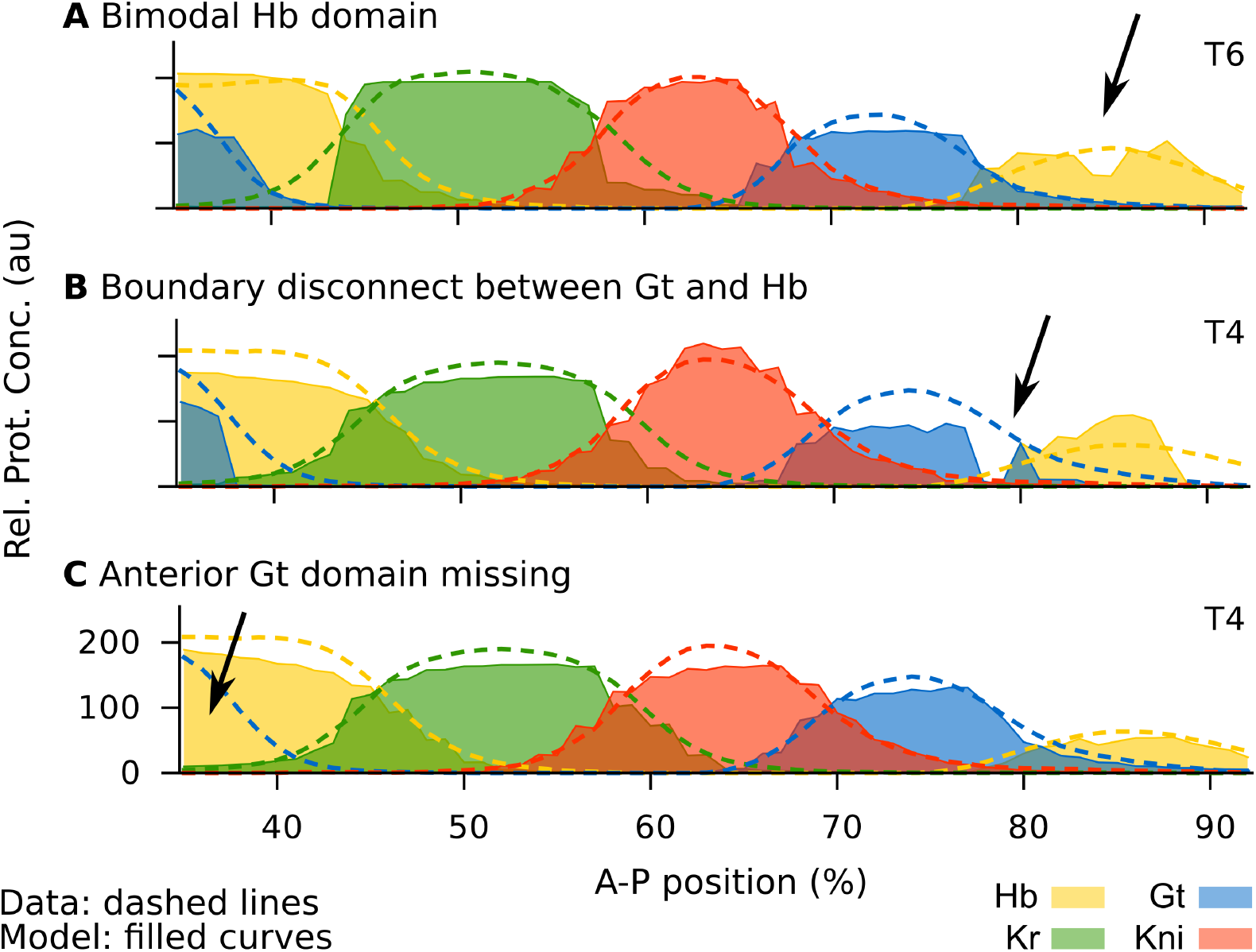
The three most commonly observed patterning defects in fully non-autonomous diffusion-less gap gene circuits. Commonly observed defects in fully autonomous *D. melanogaster* gap gene circuits fitted to data without diffusion. Circuits showing any of these gross patterning defects were excluded from further analysis, even if their RMS score was low. Arrows indicate patterning defects as named in the panel headings **(A–C)**. Horizontal axes represent %A–P position (where 0% is the anterior pole). Vertical axes show relative protein expression levels (Rel. Prot. Expr.) in arbitrary units (au). T4/6 indicate time classes C14-T4 and T6, respectively.

## Acknowledgments

We would like to acknowledge Nick Monk for countless discussions and help with the analysis of non-autonomous gene circuits. We thank Anna Kicheva and Erik Clark for critical reading and comments that significantly improved our initial drafts of this paper. We would also like to thank Manu, Lena Panok and Vitaly Gursky for their help during the early stages of this project.

